# Cold Atmospheric Plasma Differentially Affects Cell Renewal and Differentiation of Stem Cells and Apc-Deficient-Derived Tumor Cells in Intestinal Organoids

**DOI:** 10.1101/2021.10.13.464287

**Authors:** Alia Hadefi, Morgane Leprovots, Max Thulliez, Orianne Bastin, Anne Lefort, Frédérick Libert, Antoine Nonclercq, Alain Delchambre, François Reniers, Jacques Devière, Marie-Isabelle Garcia

## Abstract

Cold atmospheric plasma (CAP) treatment has been proposed as a potentially innovative therapeutic tool in the biomedical field, notably for cancer due to its proposed toxic selectivity on cancer cells versus healthy cells. In the present study, we addressed the relevance of three-dimensional organoid technology to investigate the biological effects of CAP on normal epithelial stem cells and tumor cells isolated from mouse small intestine. CAP treatment exerted dose-dependent cytotoxicity on normal organoids and induced major transcriptomic changes associated with global response to oxidative stress, fetal-like regeneration reprogramming and apoptosis-mediated cell death. Moreover, we explored the potential selectivity of CAP on tumor-like Apc-deficient versus normal organoids in the same genetic background. Unexpectedly, tumor organoids exhibited higher resistance to CAP treatment, correlating with higher antioxidant activity at baseline as compared to normal organoids. This pilot study suggests that the *ex vivo* culture system could be a relevant alternative model to further investigate translational medical applications of CAP technology.

## INTRODUCTION

Cold atmospheric plasma (CAP) is a partially or totally ionized gas including photons, electromagnetic fields, electrons, ions and neutral radicals such as reactive oxygen and nitrogen species (RONS)(1). In the past decade, this innovative technology has generated growing interest for various applications in the biomedical field such as blood coagulation, sterilization, wound healing and anti-cancer therapy(2). Number of studies have reported that CAP exerts cytotoxic effects when applied directly to cultured cells or indirectly through a plasma activated medium (PAM), as well as anti-tumoral activity(3)(4). The cytotoxic effects are mainly attributed to the production of short and long-lived RONS that generate a redox imbalance, leading to increased intracellular oxidative stress along with damage of cellular components, such as proteins, lipids and DNA(5)(6). Metabolically active cancer cells are reported to exhibit higher basal level of oxidative stress as compared to healthy cells, which could explain the reported selectivity of the anti-tumoral effects of CAP (7)(8). However, the underlying molecular mechanisms of CAP tumor selectivity remain to be clarified. Additionally, most *in vivo* studies have focused on immortalized cells, which do not recapitulate the physiological complexity of epithelia, constituted of various cell types devoted to specific functions *in vivo*(7). Lastly, the impact of CAP treatment on healthy tissues that naturally have a high renewal rate has not been fully investigated.

The adult intestinal epithelium is one of the most rapidly self-renewing tissues in adult mammals, supported by a pool of Lgr5 intestinal stem cells (ISCs), also called crypt base columnar cells, that reconstitute the whole epithelium in less than 5 days(9). ISCs have the capacity to both self-renew and give rise to transit-amplifying cells which differentiate along the villus architecture into all the cell lineages of the epithelium, (i.e., absorptive enterocytes, mucus-producing goblet cells, hormone-secreting enteroendocrine cells, Paneth cells generating antimicrobial products, and chemosensory type 2 immune response-induced tuft cells)(10). The *ex vivo* culture technology has recently been developed to indefinitely grow ISCs in a Petri dish. Upon seeding into a 3D matrix, ISCs self-renew, proliferate and differentiate into the various epithelial lineages present in the normal epithelium; *ex vivo* grown organoids maintain the *in vivo* relative proportion of each cell subtype and its temporal differentiation (11). Recently, this versatile technology has been used in the context of human colorectal(12)(13) and rectal cancer (14) allowing for the accurate prediction of drug responses(15)(16) in a personalized treatment setting.

In the present study, using the *ex vivo* culture system, we investigated the impact of an endoscopic helium plasma jet application on mouse ISCs at the morphological, cellular and transcriptomic levels. Moreover, we explored the potential selectivity of CAP application on tumor versus normal organoids originating from the same genetic background. Our data suggest that the *ex vivo* culture system could be a relevant alternative model to further investigate translational medical applications of CAP.

## RESULTS

### CAP treatment affects normal growth of intestinal stem cell-derived organoid cultures

To assess the potential benefits and adverse consequences of CAP treatment on normal surrounding tissues in the digestive epithelium, we set out to investigate the biological effects of plasma on mouse ISCs using the plasma jet device as described in Fig. 1a. For this purpose, we first generated mouse intestinal organoid lines from individual mice (Fig. 1b). For the experiments, fully grown organoids were mechanically dissociated and replated in a tridimensional matrix (Matrigel) supplemented with complete culture medium (Fig. 1b). Twenty-four hours after organoid replating (at day 1), CAP was directly applied to the culture plate wells. Different settings were applied, with varying durations (30 s or 60 s) and powers (30 W, 60 W or 80 W). Fresh culture medium was added 24 hours after CAP treatment (at day 2) and at day 4. At the endpoint (day 5), organoid survival and morphology of the grown elements were compared (Fig. 1c, d). Control cultures, and those treated with either helium gas alone (designated “60 s/0 W”) or mild CAP dose (60 s/30 W) followed similar organoid morphological evolution and survival rates. The spheroid-like structures (primarily constituted of proliferating ISCs during the first days of culture), started protruding as crypt-like domains and then differentiated into the various epithelial lineages present in the normal epithelium by day 5. Then, differentiated cells accumulated in the lumen of fully grown organoids (Fig. 1c, d). Conversely, exposure to moderate and higher doses of CAP (60 and 80 W, respectively) significantly reduced organoid budding capacity. Meanwhile, spheroid-like elements were overrepresented as compared to controls at day 5 (Fig. 1c, d). Moreover, under the 80 W-dose condition, dark declining structures were observed, accompanied by overall reduced survival rate tendency as compared to controls [34.2 ± 1.6% in CAP 60 s/80 W versus (vs) 44.5 ± 4.3% in untreated samples, respectively, unpaired t-test p=0.0698]. Together, these experiments indicate that direct CAP treatment alters the growth capacity of healthy ISCs in a dose-dependent manner.

**Figure 1.**
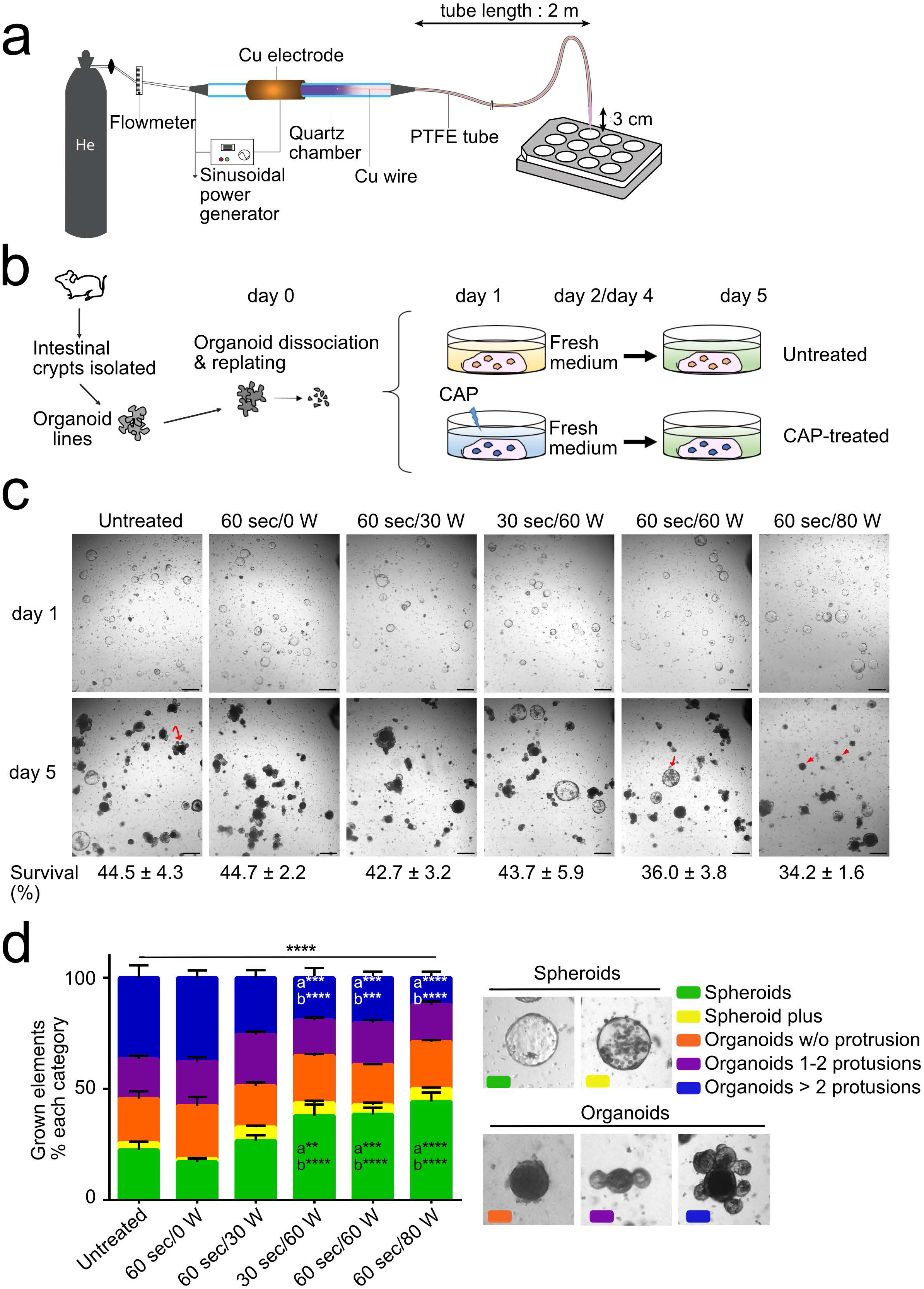
CAP treatment affects normal growth of intestinal stem cell-derived organoid cultures. **a**. The CAP system allows for generation of a helium CAP that can be transported over long distances for applications in endoscopy. **b**. Direct CAP treatment on organoid cultures generated from small intestine isolated crypts. **c**. Representative pictures of a given field showing growth of CAP-treated or untreated organoids at day 1 (before CAP application) and day 5 (endpoint). Survival rate (%) at day 5 vs day 1, indicated below the images, is expressed as the mean ± SEM (n=4 organoid lines generated from different mice). Curved and straight arrows show fully protruded organoid and spheroid, respectively. Arrowheads indicate declining organoids. Scale bars: 500 μm. **d**. Quantification of organoid complexity at day 5. Representative element categories (Spheroids and Organoids) are defined morphologically on the right. An average number of 195 elements was analyzed per condition per organoid line (n= 3 organoid lines). Data are represented as means ± sem. Two-way ANOVA: interaction **** P< 0.0001 followed by Tukey’s multiple comparisons test: a: compared to Untreated conditions, b: compared to 60 s/ 0 W conditions **P< 0.01; ***P< 0.001; ****P< 0.0001.

Next, we investigated the influence of the culture medium on ISC growth at a fixed moderate dose (60 s/ 50 W) (see experimental design in Fig. 2a). Direct CAP treatment was applied at day 1 and the treated culture medium was replaced with fresh medium at 30 min, 3 h or 1 day (1 d) later (Fig. 2a). The morphology of grown elements was studied at day 3, an endpoint at which 61.8% of grown elements in untreated cultures showed at least one crypt-like domain (Fig. 2b, c). Removal of CAP-treated medium early after the procedure (30 min and 3 h) did not significantly alter protrusion formation. In contrast, prolonged incubation with CAP-treated culture medium (24 hours = 1 d) was associated with higher proportion of elements grown as spheroids (67%), only 9% of total elements were protruded organoids (Fig. 2b, c). Together, these data indicate that the time of exposure to reactive species generated by CAP treatment affects the capacity of ISCs to normally grow and differentiate. Interestingly, a similar negative impact on organoid budding was observed when “naïve” organoids were indirectly submitted to reactive species present in conditioned media generated either by CAP treatment applied onto another organoid culture (designated as “indirect CAP”) or the Plasma-Activated Medium alone (designated as “PAM”) (Fig. 2a-c). Exposure to Indirect CAP- or PAM-conditioned media 24 hours after the treatment (from day 2 to day 3) also altered organoid protrusion, albeit to a milder degree, thereby indicating that toxic long-lived reactive species were still present in culture supernatants after 24 hours (Supplementary Fig. 1S a-c). Taken together, these data demonstrated that normal ISCs are sensitive to direct as well as indirect CAP/PAM treatment *ex vivo*.

**Figure 2.**
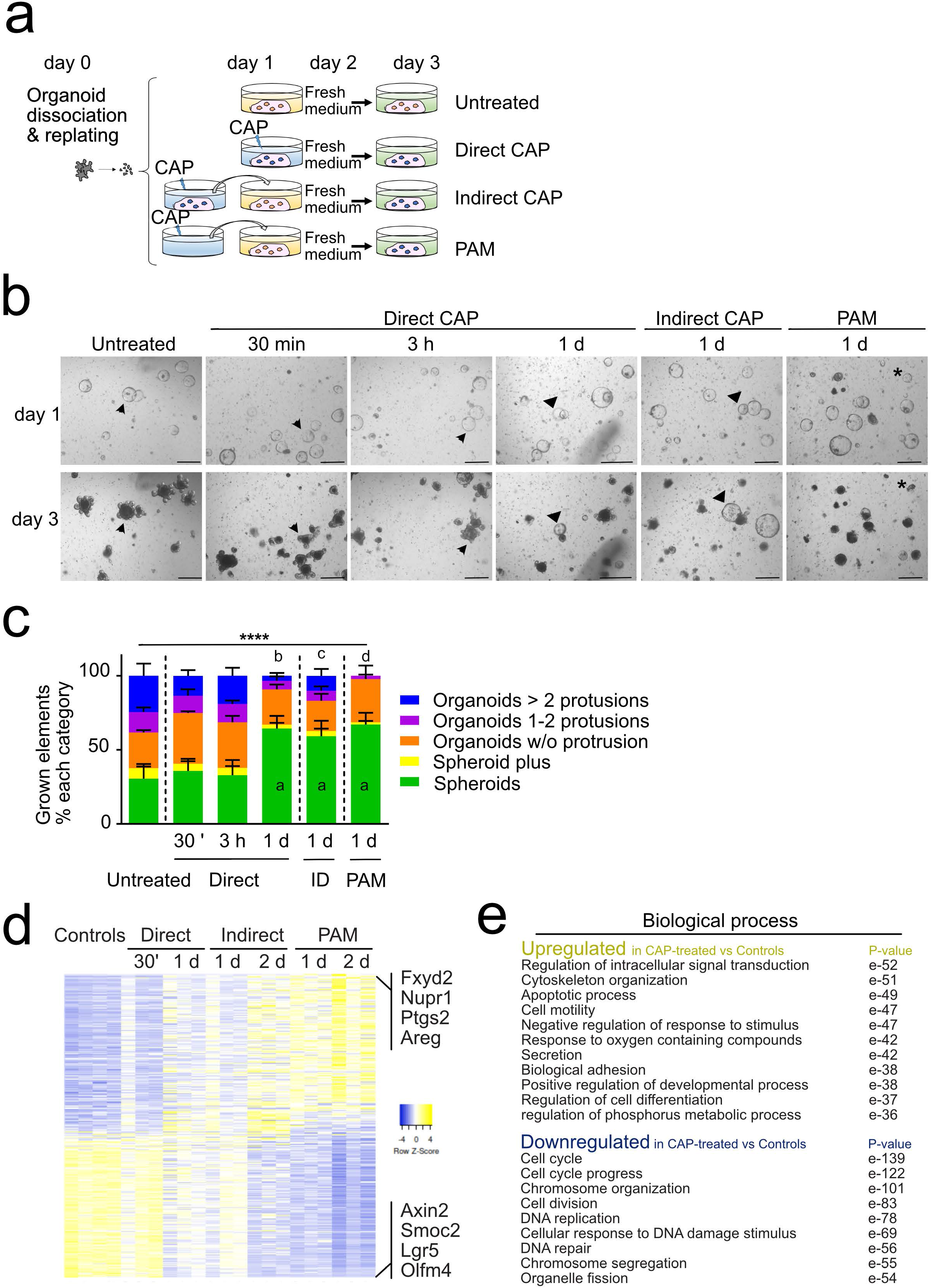
Impact of the CAP application method on organoid morphology and global gene expression. **a**. CAP treatment (50 W/60 s) was applied directly to organoid cultures at day 1 (post-replating) for 30 minutes, 3 hours or 24 hours. Then fresh medium was provided until day 3 (endpoint). Alternatively, naïve organoids were cultured at day 1 for 24 hours with freshly-generated CAP-conditioned media (Indirect CAP) or plasma-activated media (PAM). At day 2 (post-replating), fresh culture medium was provided for a further 24 hours until day 3 (endpoint). **b**. Representative pictures of a given field showing organoid growth at day 1 (before CAP application) and day 3. Arrowheads and triangles show individual elements evolving as protruded organoids and spheroids, respectively. Asterisks indicate dying elements. Scale bars: 500 μm. **c**. Quantification of organoid complexity at day 3. An average number of 100 elements was analyzed per condition per organoid line (n= 4 organoid lines). Data are represented as means ± sem. Two-way ANOVA: interaction **** P< 0.0001 followed by Tukey’s multiple comparisons test: a, d: ****P< 0.0001, b: ***P< 0.001, c: *P< 0.05. (all compared to untreated). Time of exposure to CAP treatment is indicated (30’, 3 h, 1 day) ID: Indirect CAP. **d**. Heatmap of differentially regulated genes in CAP-treated vs Untreated (Controls) organoids. Samples were treated with CAP directly (Direct), for 30 min (30’) or 24 hours between day 1 and day 2 (1 d); indirectly (Indirect) or with PAM for 24 hours between day 1 and day 2 (1 d) or between day 2 and day 3 (2 d). Some of the most-modulated genes are indicated on the right side. **e**. GSEA-Biological processes for upregulated and downregulated gene lists in CAP-treated vs untreated Controls.

### CAP treatment decreases the intestinal stem cell pool and is associated with Apoptosis

To further investigate the molecular and cellular mechanisms of CAP-induced morphological effect on intestinal organoid growth, we analyzed at day 3 the whole transcriptome of organoids treated directly or indirectly with CAP at a moderate dose (50W/60 s) during a period of 24 hours (Fig. 2a). We identified 2462 differentially expressed genes in CAP-treated vs untreated cultures (False Discovery rate 0.01 and Log2-fold change of 1 or above) (Fig 2d). Of these, CAP-treatment induced downregulation of 1195 genes involved in the following biological processes: cell cycle progression, cell division and DNA replication (Fig. 2e). ISC markers like *Lgr5*, *Olfm4*, *Smoc2* and *Axin2* genes were amongst the most downregulated genes (Fig. 2d). In particular, 35% of the genes identifying the intestinal crypt base columnar stem cell signature (i. e. 135 out of 379 genes) reported by Munoz et al, (17) were downregulated in CAP-organoids (Fig. 3a, b). Loss of ISCs in CAP-treated cultures was confirmed by *in situ* hybridization experiments and immunofluorescence staining for *Olfm4*-expressing cells; this correlated with significant drop in canonical Wnt signaling activity (*Axin2* used as reporter gene) (Fig. 3b-d). Consistent with suggested upregulation of the apoptotic process (p value e-49, Fig. 2e), the Pycard gene, involved in regulation of apoptosis and adaptation to the inflammatory response, and genes associated with DNA repair (*Pclaf*, *Usp49*, *Hmga2*) were found to be downregulated meanwhile apoptosis-mediator/effector genes (Calpains *Capn2/5/12/13*, *Trp53inp2*, *Ctsl*, *Unc5b*, *Cidec*, *Atf3*, *Ddit3*, *Cfap157*) were upregulated in CAP-treated organoids (Fig. 3c, 4a). TUNEL assay performed on organoid sections confirmed the presence of DNA double strand breaks in CAP-treated cultures (Fig. 4b).

**Figure 3.**
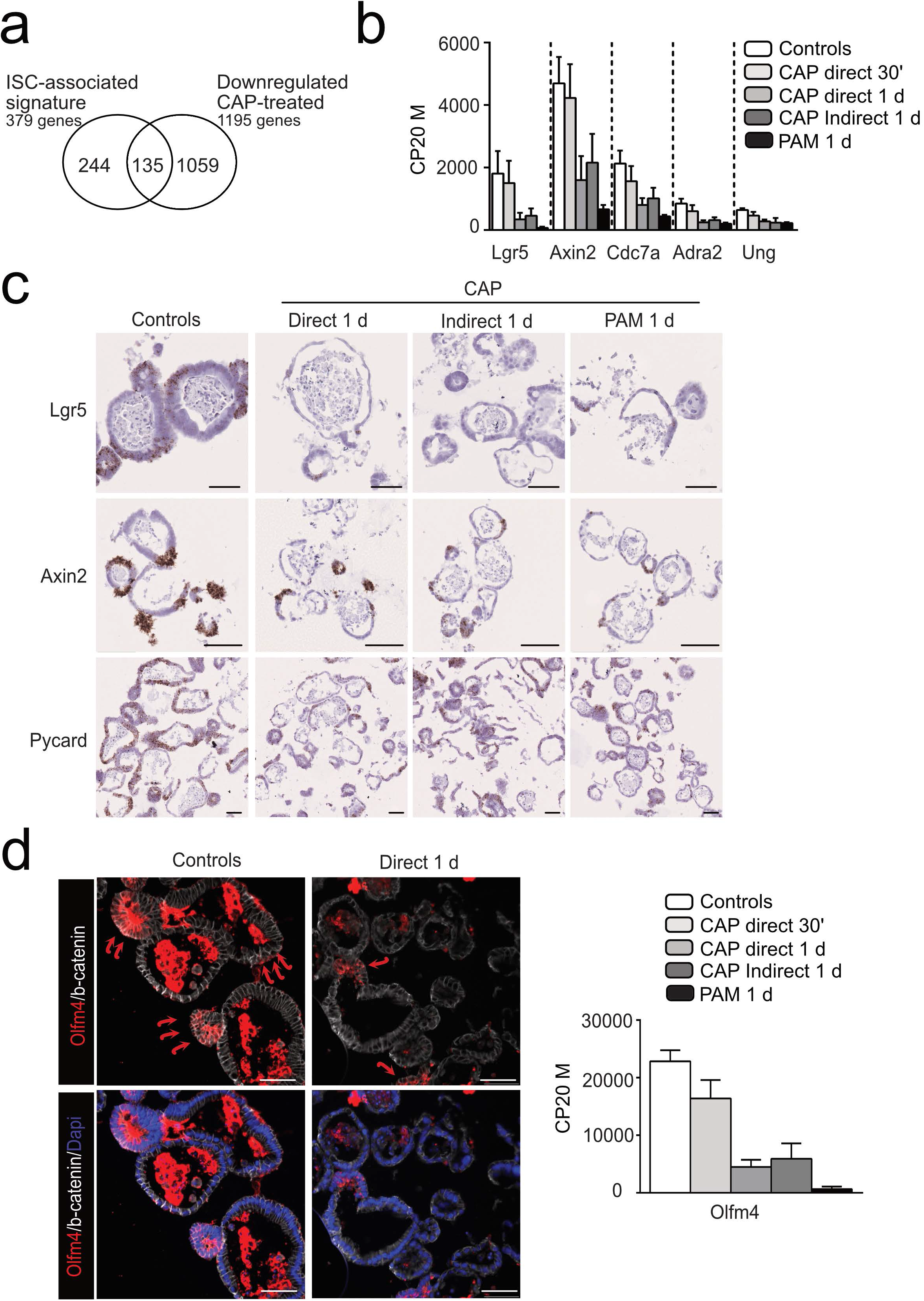
CAP treatment decreases the intestinal stem cell pool. **a**. Venn diagram showing that part of the downregulated gene list in CAP-treated samples vs controls (135 out of 1195 genes) corresponds to the ISC (Intestinal Stem Cell)-associated signature. **b**. Expression levels of some genes of the ISC-associated signature commonly downregulated in CAP-treated organoids vs Untreated controls. CP20M: counts per kilobase of transcript per 20 million mapped reads. Data are represented as means ± sd. n = 4 and 3 samples in Controls and CAP-treated conditions, respectively. **c**. Expression of *Lgr5*, *Axin2,* and *Pycard* genes detected in organoids by RNAscope at day 3. **d**. Immunofluorescence showing Olfm4-expressing cells in organoids at day 3 (red arrows). Cell membranes shown with β-catenin and nuclei counterstained with DAPI. Right panel: Expression levels of *Olfm4* in the various conditions reported in CP20M. Scale bars: 50 μm (panels c and d).

**Figure 4.**
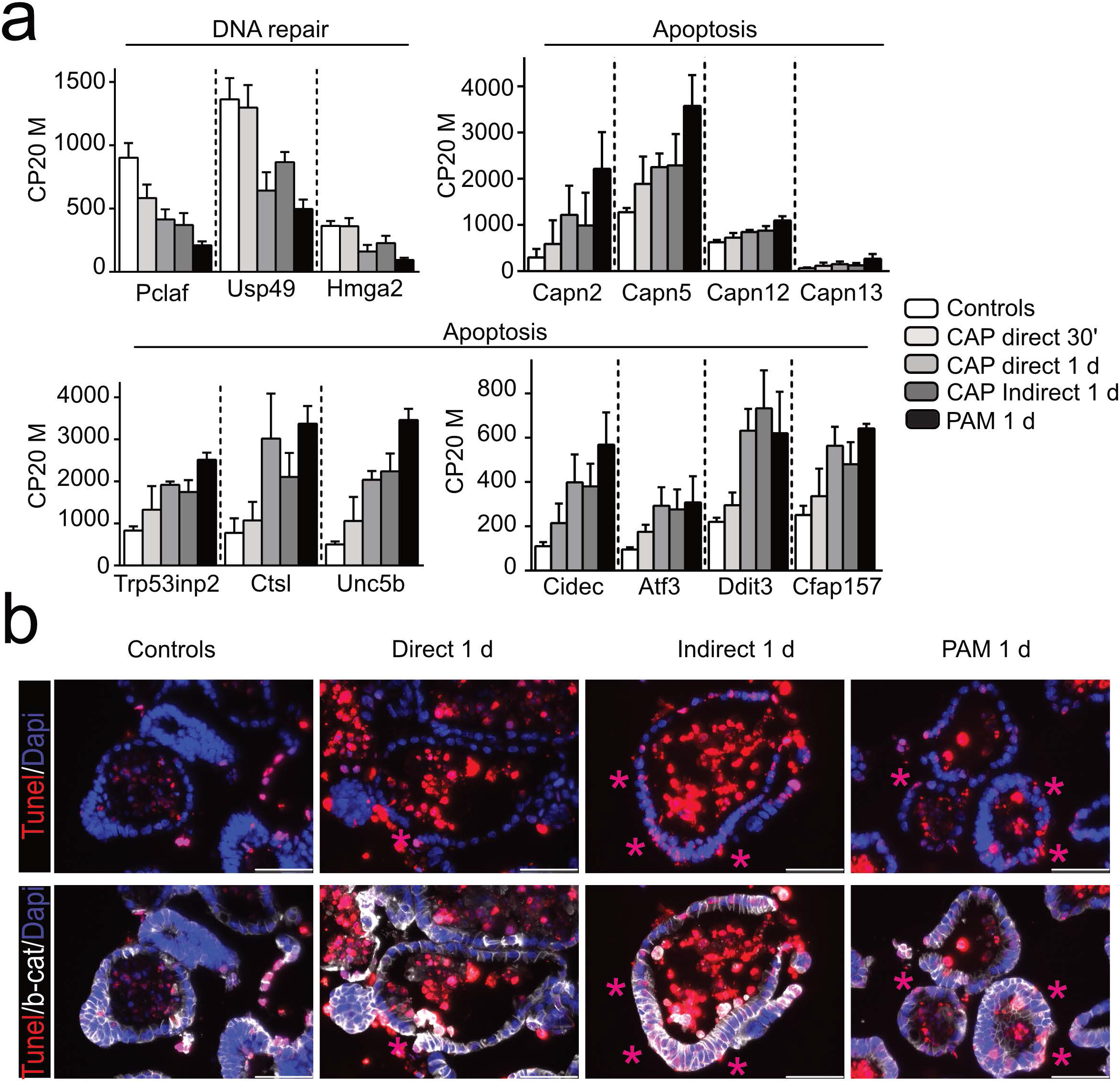
CAP treatment of organoids is associated with Apoptosis. **a**. Expression levels of DNA repair and apoptosis-related genes in the various conditions reported in CP20M. Data are represented as means ± sd. n = 4 and 3 samples in Controls and CAP-treated conditions, respectively. **b**. Organoid sections were stained with TUNEL for apoptotic cells (visualized by pink asterisks). Cell membranes are shown with β-catenin and nuclei were counterstained with DAPI. Scale bars: 50 μm.

### CAP treatment induces a global response to reactive species in intestinal stem cell-derived organoids

In addition, CAP induced upregulation of genes involved in modulation of intracellular signal transduction, negative regulation of response to stimulus, and response to oxygen containing compounds as well as in positive regulation of developmental processes (Fig. 2e). In line with response to CAP mediated-injury, pro-survival (*Dsg3*, *Phlda3*) and regeneration-associated genes (*Trop2*/*Tacstd2*, *Ptgs2*, *Hbegf*, *Ly6a*/*Sca1*, *Clu*, *Areg*, *Epn3*) were induced as early as 30 min post-direct CAP treatment (Fig. 5a). Indeed, 34% of the genes identifying the fetal-like Trop2 regeneration signature (50 out of the 148 gene list, (18)) were upregulated in CAP-treated organoids (Fig. 5b). Accordingly, both the number of Trop2-expressing cells and Areg expression were significantly increased in CAP-treated organoids (Fig. 5c, d). Furthermore, a global stress response gene expression pattern was detected with early upregulation of oxidative stress-associated transcription factors like *Fos*, *Fosb*, *Egr1*, *Nfe2l1*/*Nrf1*, *Nupr1*, *Nr4a1* and *Jdp2* (Fig. 5a, Supplementary Fig. 2Sa). In line with reported regulation of oxidative stress by long noncoding RNAs, expression of *Fer1l4*, *Gm20417*, *Neat1*, *Malat1*, *Gm37376* or *Kcnq1ot1* was induced 30 min after CAP-treatment (Supplementary Fig. 2Sa) (19). Effectors of global response to CAP-mediated oxidative stress in organoids were found upregulated. Among these, genes encoding solute/metabolite transporters such as *Slc7a11*, a cysteine/glutamate antiporter xCT, and detoxifying enzymes (*Hmox1*, *Ethe1*, *Gch1*, *Gstk1*) were identified (Fig. 5a,d). Components of the cysteamine/cystamine metabolism (*Gpr5a*, *Vnn1*, *Chac1*, *Ggh*), involved in glutathione redox status, were also induced by the treatment (Supplementary Fig. 2Sb). Cell signaling molecules such as the Polo like kinases (*Plk2*, *Plk3*) reported to exert antioxidant functions, as well as *Mapk11*, *Lif* and *Pmepa1* were also upregulated by CAP application (Fig. 2Sb). Of note, the major cellular enzymatic ROS scavengers (superoxide dismutases, glutathione peroxidases, peroxiredoxins and catalase) were detected at high levels in control organoids but their expression did not substantially differ upon CAP-treatment (Table 1). *Pparg*, a master regulator of lipid metabolism and immune response (20), was also found upregulated in CAP-treated organoid cells (Fig. 5a, d). Inflammatory response genes were significantly modulated in CAP-treated organoids: interferon induced *Ifitm2/Ifitm3* and chemokine *Ccl9* were downregulated whereas expression of *Ifnlr1, Ilr1, Ddx60* and *Pdlim7* were upregulated (Fig. 2Sb). Moreover, consistent with extensive reshaping of epithelial cells, CAP application was associated with upregulation of cytoskeleton organization, cell motility and biological adhesion processes (Fig. 2e, Fig. 2Sb).

**Table 1:**
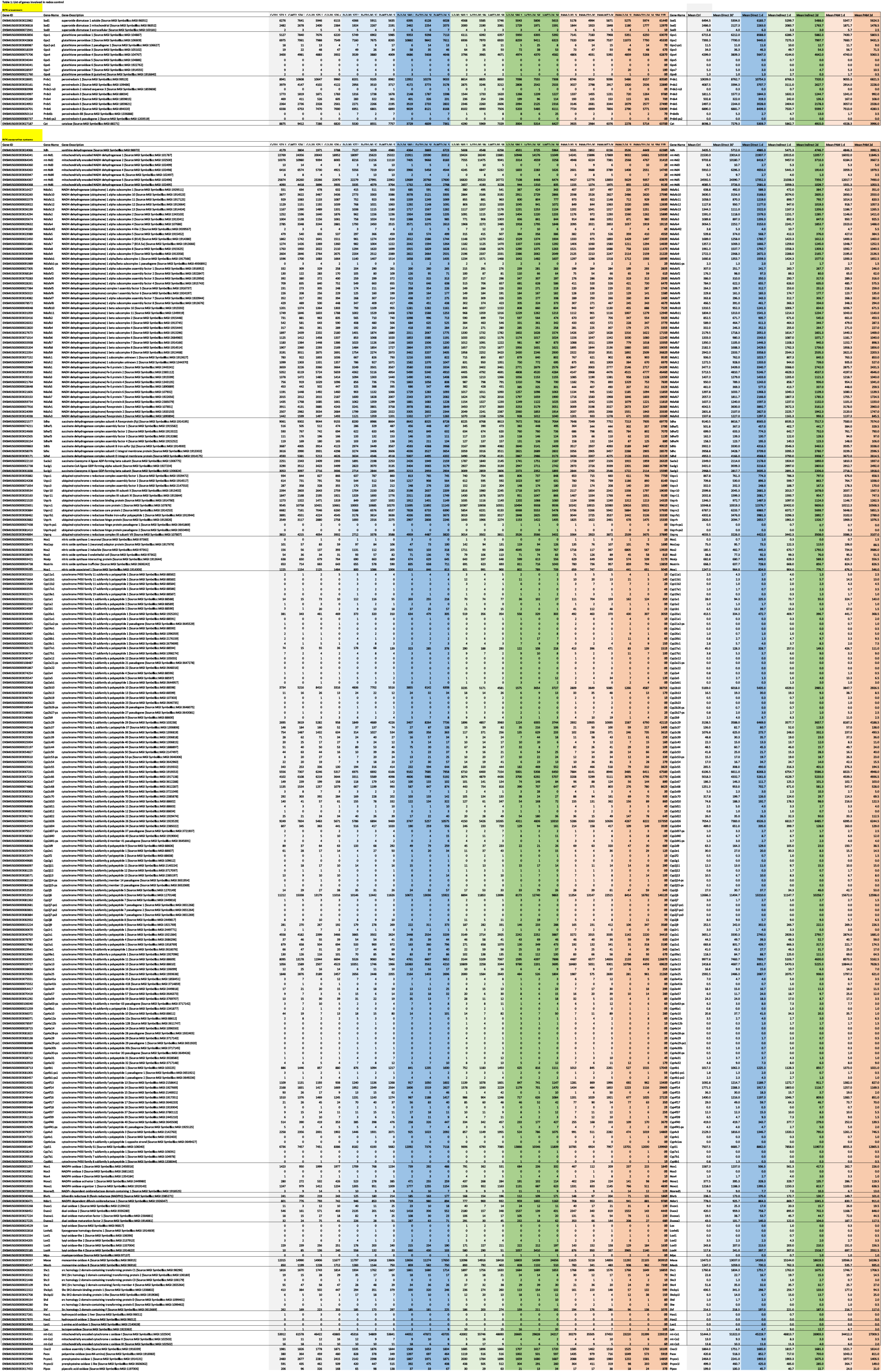
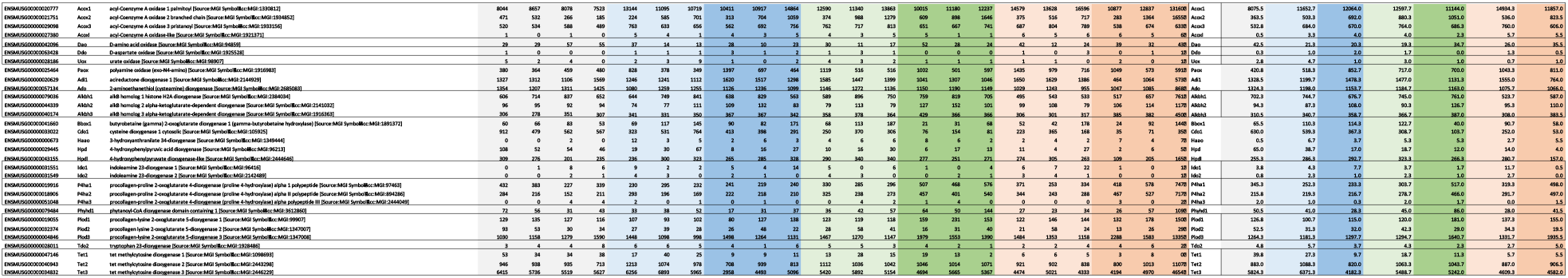
List of genes involved in redox control

**Figure 5.**
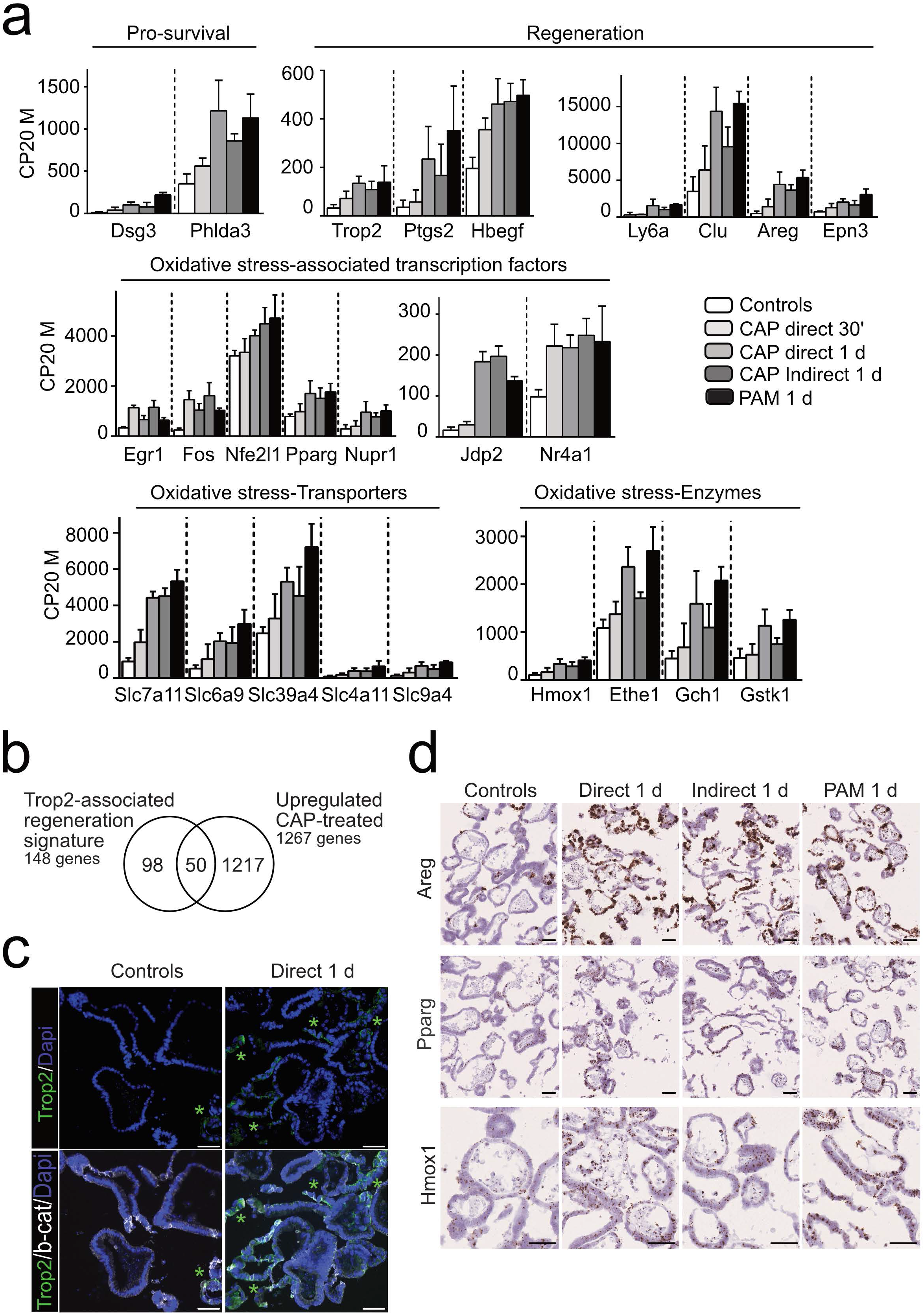
CAP treatment induces a global response to reactive species in intestinal stem cell-derived organoids. **a**. Expression levels of Pro-survival, regeneration and oxidative stress-associated genes in the various conditions reported in CP20M. Data are represented as means ± sd. n = 4 and 3 samples in Controls and CAP-treated conditions, respectively. **b**. Venn diagram showing that part of the upregulated gene list in CAP-treated vs control organoids (50 out of 1267 genes) corresponds to the Trop-2-associated regeneration signature. **c**. Immunofluorescence showing Trop2-expressing cells (visualized by green asterisks) in organoids at day 3. Cell membranes shown with β-catenin and nuclei counterstained with DAPI. **d**. Expression of *Areg*, *Pparg,* and *Hmox1* genes detected in organoids by RNAscope at day 3. Scale bars: 50 μm (panels c and d).

### Apc deficient-derived organoids exhibit increased resistance to CAP treatment as compared to ISCs-derived organoids

Since ISCs were sensitive to moderate and high doses of CAP, we sought to compare resistance of normal or tumor organoids to this treatment. For this purpose, adult VilCreERT2-Apc^*flox/flox*^ or VilCreERT2-Apc wild-type (wt) mice were injected with tamoxifen to induce specific deletion of Apc exon 15 in VilCreERT2-Apc^*flox/flox*^ mice, leading to loss-of-function of this tumor suppressor (designated as Apc Δ). Apc wt and Apc Δ intestinal crypts were isolated and cultured to generate normal and tumor organoid lines, respectively (Fig. 6 a). Efficient recombination in Apc Δ tumor-derived organoids was controlled at initial seeding (Supplementary Fig. 3Sa). Then, upon organoid replating, CAP was directly applied at various doses for 60 s on both kinds of organoids (Fig. 6b). As expected, at day 5, Apc wt organoid survival was substantially reduced at a dose of 50 W and 80 W as compared to untreated organoids (Fig. 6 b,c). Conversely, Apc Δ organoids demonstrated higher survival rates, regardless of the CAP dose (Fig. 6c). Dosage of reactive species in culture supernatants did not demonstrate substantial differences between organoid types at a given CAP-dose, which might have explained the observed higher resistance of Apc Δ vs Apc wt organoids (Supplementary Fig. 3Sb). To further explore the underlying molecular mechanisms, qPCR experiments were performed on samples collected at day 5. With the exception of *Olfm4*, expression of markers for active (Lgr5) and quiescent (Hopx) stem cell populations was maintained in tumor organoids, even at the highest CAP dose (80 W) (Fig. 6d). Regarding cell differentiation, mild CAP dose (30 W) in Apc wt organoids had a tendency to decrease the Paneth lineage vs the other cell types whereas Apc Δ-organoids exhibited limited cell differentiation in any tested condition (Fig. 6d). Interestingly, expression of regeneration marker genes was upregulated in Apc wt organoids upon CAP treatment (30 W), and was detected at much higher levels in tumor organoids even under normal conditions (Fig. 7b). Moreover, tumor organoids maintained proliferation capacity and low cell death behavior following CAP treatment (Fig. 7b). Furthermore, Apc Δ organoids demonstrated increased levels of genes (e. g. *Atf3*, *Nupr1*) involved in early response to reactive species as compared to Apc wt organoids (Fig. 7b). In addition, cellular transporters (*Slc7a11*) and detoxifying enzymes (*Hmox1*), induced in Apc wt organoids by mild CAP treatment, were expressed at significantly higher levels in basal conditions in tumor organoids (Fig. 7b). Differential expression of the membrane-associated aquaporins has also been proposed to contribute to CAP selectivity by facilitating influx of water and hydrogen peroxide into cancer cells(21). In ISCs and healthy intestinal organoids, RNAseq data showed that *Aqp1*, *Aqp4 and Aqp11* were the most significantly expressed (Supplementary Fig S3c). CAP treatment particularly reduced *Aqp1* and *Aqp4* levels in Apc wt organoids (Supplementary Fig S3c). Interestingly, irrespective of CAP treatment, tumor organoids were expressing 2-fold less *Aqp1*, *Aqp4* and 10-fold less *Aqp3* levels than normal organoids, whereas Aqp5 expression was increased (Supplementary Fig S3d). Taken together, these experiments indicate that tumor-like organoids deficient for the tumor suppressor Apc demonstrate a higher potential to resist CAP-induced injury as compared to ISC-derived organoids. Such behavior could be, in part, explained by the expression of an oxidative stress resistance program, already active under basal conditions, and reduced cell surface expression of aquaporins.

**Figure 6.**
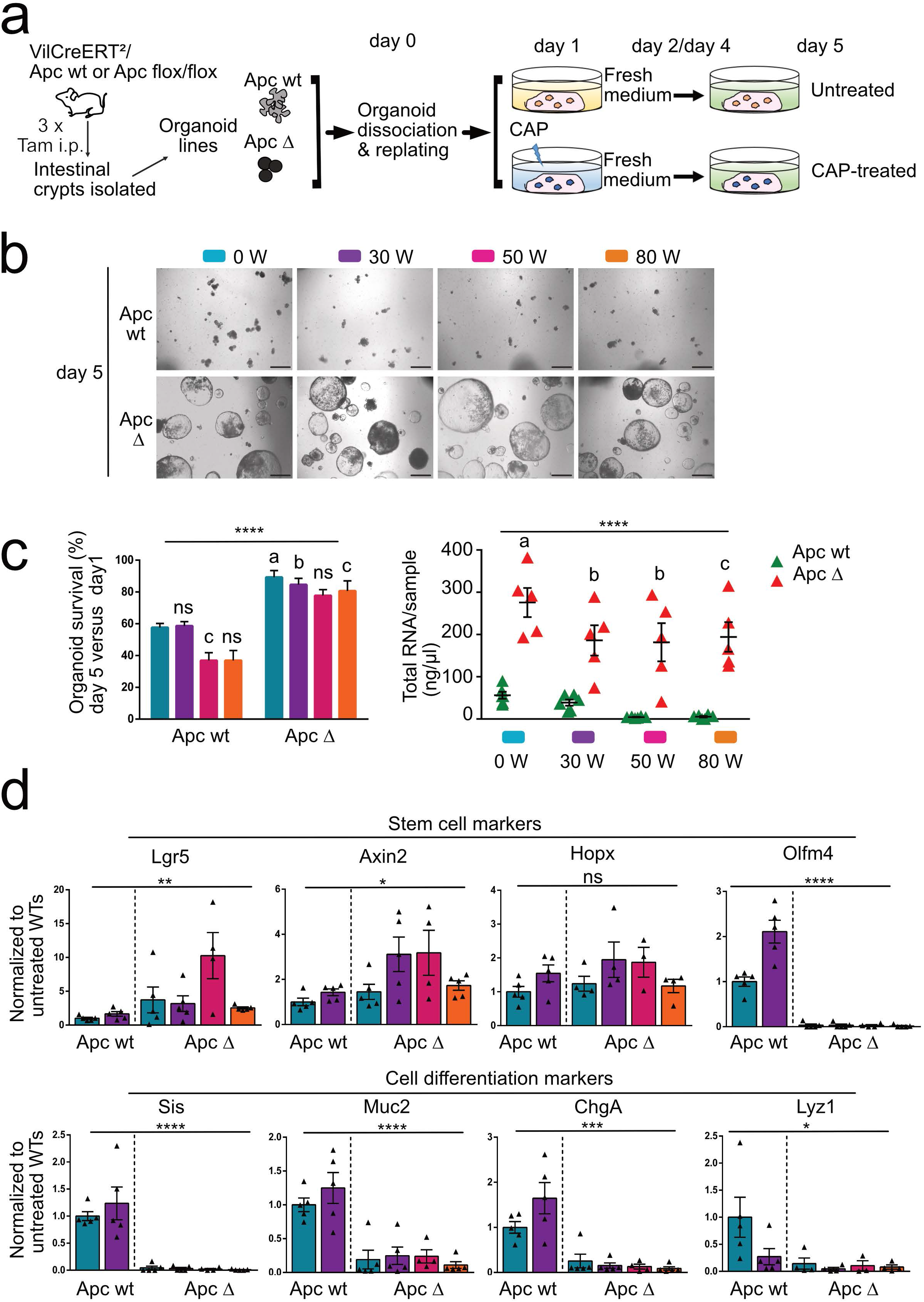
Apc deficient-derived organoids exhibit increased resistance to CAP treatment as compared to normal ISC-derived organoids. **a**. CAP treatment was applied directly to Apc wt or Apc Δ (Apc-deficient) organoid cultures at day 1 post-replating for 24 hours. Fresh medium was added at days 2 and 4. **b**. Representative pictures of a given organoid type (Apc wt or Apc Δ) at day 5 following CAP treatment at the indicated doses. Scale bars: 500 μm. **c**. Left panel: organoid survival rate (in %). An average number of 120 elements was studied over time per condition per organoid line (n= 6 Apc wt and 5 Apc Δ organoid lines, respectively). Data are represented as means ± sem. Two-way ANOVA: CAP dose ***/Genotype **** P< 0.0001 followed by Sidak’s multiple comparisons test: a, ***P< 0.001; b, ***P< 0.01; c, *P< 0.05; ns: not significant (all compared to Apc wt-0 W), Right panel: quantification of total RNA extracted/sample at day 5. Each symbol corresponds to a given organoid line. Two-way ANOVA: CAP dose * P< 0.05/Genotype **** P< 0.0001 followed by Sidak’s multiple comparisons test: a, ****P< 0.0001; b, *P< 0.05; c, **P< 0.01. (all compared to Apc wt-0 W). **d**. Gene expression analysis by qRT-PCR of the indicated stem cell and differentiation markers. Each symbol corresponds to a given organoid line. Values are normalized to Untreated Apc wt levels. Data are represented as means ± sem. One-way Anova test. **** P< 0.0001; **** P< 0.001; **P< 0.01; *P< 0.05; ns: not significant.

**Figure 7.**
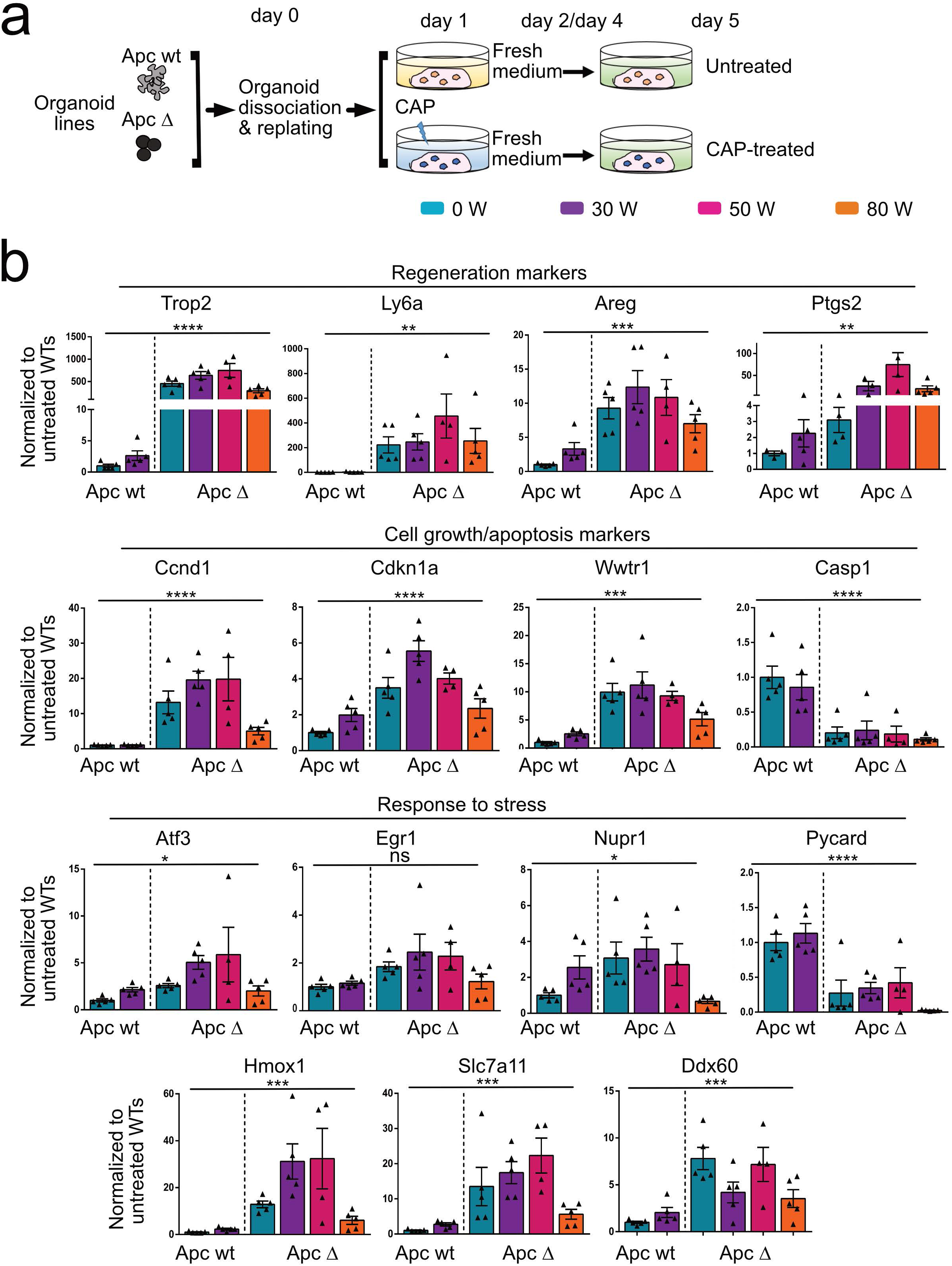
Apc deficient-derived organoids exhibit increased resistance to CAP treatment as compared to normal intestinal stem cell-derived organoids. **a**. Design of the experiment. See Figure Legend Fig. 6a. **b**. Gene expression analysis by qRT-PCR of the indicated markers involved in regeneration, cell growth/apoptosis and response to stress. Each symbol corresponds to a given organoid line. Values are normalized to Untreated Apc wt levels. Data are represented as means ± sem. One-way Anova test. **** P< 0.0001; **** P< 0.001; **P< 0.01; P< 0.05; ns: not significant.

## DISCUSSION

The present study demonstrated the relevance of a 3D organoid culture to investigate the biological effects of CAP on normal epithelial stem cells and tumor cells isolated from the mouse small intestine, and unexpectedly showed higher resistance of tumor organoids to CAP treatment.

First, we studied the parameters that determine CAP effects on cells and found that helium, the carrier gas, did not by itself alter organoid growth. We observed toxic effect of direct CAP application to ISC cultures, showing dose-dependency as reported on various cancer lines (21). To determine whether CAP treatment can induce a “bystander effect” as described in radiobiology, in which irradiated cells communicate stress via extracellular signaling to neighboring cells, we also analyzed indirect application of CAP on organoids (Indirect CAP and PAM). In our experimental setting, organoid growth was similarly affected by the three conditions, indicating that the main effect of CAP was mediated by reactive species delivered in the culture medium, with minor contribution of cellular components potentially released from the CAP-treated organoids. Our data also suggest that CAP-mediated cytotoxicity depends in part on long-lived reactive species (typically nitrites, hydrogen peroxide, ozone) since increased levels of nitrites were still detected 24 hours after CAP treatment as compared to controls (Supplementary Fig S3b). On the other hand, it is known that the nature of CAP-generated RONS is conditioned by the composition of the cell culture medium(22). Organoid growth requires a complex medium, including several antioxidant supplements such as N-acetyl cysteine, glutathione, sodium pyruvate and ascorbic acid (23) (24)(25) (26). We hypothesize that the presence of such antioxidants in the medium might have increased the threshold level needed to detect variations in the long-lived ROS species with the DCFH-DA probe (Supplementary Fig S3b).

Moderate (50/60 W) to high (80 W) doses of CAP severely impacted ISC self-renewal and differentiation. Although ISCs express important levels of ROS scavenger enzymes (Supplementary Fig. S3e), previous studies have also reported that, under steady-state conditions, ISCs exhibit two-fold increased levels of ROS as compared to the rest of the epithelium(27). Together, these data are in line with the hypothesis that the application of moderate-to-high CAP doses to ISCs reaches a redox status threshold above which toxic accumulation of RONS occurs, similar to what has been previously described for cancer cell lines(7)(8); this ultimately induces DNA damage and cell death by apoptosis. Interestingly, moderate CAP dose also elicited an epithelial response with hallmarks of tissue regeneration, characterized by re-expression of a fetal signature, reported to be induced by a variety of insults in the gastrointestinal tract(18)(28)(29)(30). As part of this fetal reprogramming, upregulation of genes coding for factors mainly released by the surrounding stromal compartment (Areg, Ptgs2, etc..), was observed in epithelial cells *ex vivo*. This suggests that, upon injury, the epithelium has the intrinsic potential to contribute to the repair process. In the same line, CAP treatment on murine cementoblasts has been reported to induce a regeneration process similar to that elicited by enamel matrix derivatives *in vivo*(31). Moreover, CAP treatment induced a global response to oxidative stress in organoids that involved upregulation of RONS-associated transcription factors and effectors known to regulate intracellular ROS balance and reported to be activated upon CAP treatment(32)(33)(34)(35).

A major observation of our study relates to CAP application effects on tumoral APC-deficient organoids. The potential of CAP treatment in oncology relies on a proposed selective targeting of cancer cells over healthy cells due to higher intracellular levels of ROS in malignant cells (36). However, compared sensitivity of CAP was never performed on the same cell type and genetic background, with cells growing under the same culture conditions. This was addressed in the present study by comparing the impact of CAP application on normal “healthy” organoids and tumoral Apc-deficient organoids, the latter being used as a colorectal cancer (CRC) model. Contrary to our initial expectations, tumor organoids exhibited higher resistance to CAP treatment than healthy organoids. Such results could be explained, at least in part, by the fact that Apc-deficient organoids exhibited a much higher antioxidant response at baseline than ISC-derived organoids. The potential contribution of differential expression of aquaporins to CAP selectivity could be interesting to address in future *ex vivo* studies.

In summary, the present study reveals the potential of organoid technology to further investigate the biological effects of CAP on normal and tumoral tissues. Indeed, organoid cultures faithfully reflect cell heterogeneity in epithelial tissues, they are as amenable to “Omics” studies as the cancer cell lines and, from a translational point of view, they can be used to compare various CAP application settings. Our study highlights the importance of considering CAP toxicity on metabolically active resident stem cells in tissues, like the intestine, undergoing permanent self-renewal, in order to adjust the dose of treatment. Furthermore, CRC development is known to be a multistep process. Following the initial hit mutation in the Apc gene that leads to Wnt signaling overactivation, mutations deregulating other pathways (such as KRAS/TGFb and p53) sequentially accumulate in cancer cells, and correlate with cancer progression(37)(38). Future studies will be needed to investigate the sensitivity of CAP treatment on tumor organoids bearing the aforementioned additional mutations in order to definitely elucidate the potential of CAP for personalized anti-cancer therapy.

## MATERIALS AND METHODS

### Experimental animals

Adult mice were from the outbred CD1 strain (Charles River Laboratories). To obtain tumor-derived organoids, we crossed Tg(Vil1-cre/ERT2)^23Syr/J^ (39) and Apc^tm1Tyj/J^ designated Apc^flox^ (40). Adult Vil-cre/ERT2/Apc^wt/wt^ and Vil-cre/ERT2/Apc^flox/flox^ were injected intraperitoneally for 3 consecutive days with tamoxifen (2 mg per 30 g of body weight) to induce recombination in the Apc locus and the small intestine was harvested 2-3 days after the last injection. Tamoxifen was dissolved in a sunflower oil/ethanol mixture (9:1) at 10 mg/ml (both products from Sigma-Aldrich).

### Ex vivo culture

To generate intestinal organoids, mouse adult small intestine was dissociated with 5 mM EDTA-in DPBS (Gilbco) according to the protocol reported in(41). Briefly, the culture medium consisted of Advanced-DMEM/F12 medium supplemented with 2 mM L-glutamine, N2 and B27 w/o vit.A, gentamycin, penicillin-streptomycin cocktail, 10 mM HEPES (all from Invitrogen), 1 mM N acetyl cysteine (Sigma-Aldrich), 50 ng/ml EGF and 100 ng/ml Noggin (both from Peprotech), and 100 ng/ml CHO-derived mouse R-spondin 1 (R&D System). Culture medium was changed every other day and after 5-6 days in culture, organoids were harvested, mechanically dissociated and replated in fresh Matrigel matrix (catalog 356235 from BD Biosciences). Culture media were supplemented with 10 μM Y-27632 (Sigma Aldrich) in all initial seeding and replating experiments for the first two days. Pictures were acquired with a Moticam Pro camera connected to Motic AE31 microscope.

### Cold atmospheric plasma treatment

The cold plasma source was an endoscopic plasma jet allowing for the generation and transport of CAP over long distances as described in our previous work(42). It consists of a tubular DBD chamber made of quartz supplied with helium gas and surrounded by a high-voltage electrode. The power-control source was an AFS (G10S-V) generator, delivering an 18 kHz sinusoidal signal. The discharge chamber was plugged into a polytetrafluoroethylene (PTFE) tube (outer diameter 3 mm, wall thickness 0.75 mm) transporting the plasma post-discharge over > 2 meters. An electrically floating copper wire (diameter 0.2 mm) was inserted partially into the dielectric chamber and extended almost until the end of the PTFE tube (5 mm before its end) to allow the maintenance of active plasma for several meters and to sustain a plasma plume at the outlet for potential endoscopic treatment. Treatment was performed under a laminar flow hood using a 1.6 lpm helium flow; thus RONS were created by the mixing of CAP with ambient air (42). The power values (0, 30, 50, 60 or 80 W) were set on the AFS-generator. The catheter was placed vertically in a designated test bench. The tip of the catheter was placed at 3 cm from the top of the 12-well plate containing the media to be treated, i.e. the length of the plasma plume. This allowed plasma-generated RONS to reach the medium without the plasma plume touching it, avoiding any electrical connection that would have increased current and RONS creation in a less controlled manner. Fully-grown organoid cultures were dissociated by mechanical pipetting and replated on Matrigel in 12-well plates (VWR, Belgium). Twenty-four hours later, CAP was applied vertically on organoid cultures at room temperature. Each well was placed successively under the plasma plume for 30 s or 60 s. During treatment, the other wells were protected by a designated cover plate to avoid unwanted RONS diffusion to neighboring wells. Following exposure to CAP, treated samples were placed back at 37 °C in a 5% CO2 incubator (Binder C150) and the medium was changed as indicated in the results section and different endpoints were applied.

### Tissue processing and immunohistochemical analysis

Organoid culture samples were fixed with 10% formalin solution, neutral buffered (Sigma-Aldrich) for 20 minutes at room temperature then sedimented through 30% sucrose solution before OCT embedding. Histological protocols as well as immuno-fluorescence/histochemistry experiments on 6 μm sections were carried out as previously described (43). Table 2 lists primary antibodies, TUNEL assay kit and DAPI used. Stained samples were visualized with a DMI600B epifluorescence microscope equipped with a DFC365FX camera (Leica). The number of independent organoid cultures obtained from individual adult mice used for each experiment is reported in Figures and in Figure legends.

**TABLE 2.**
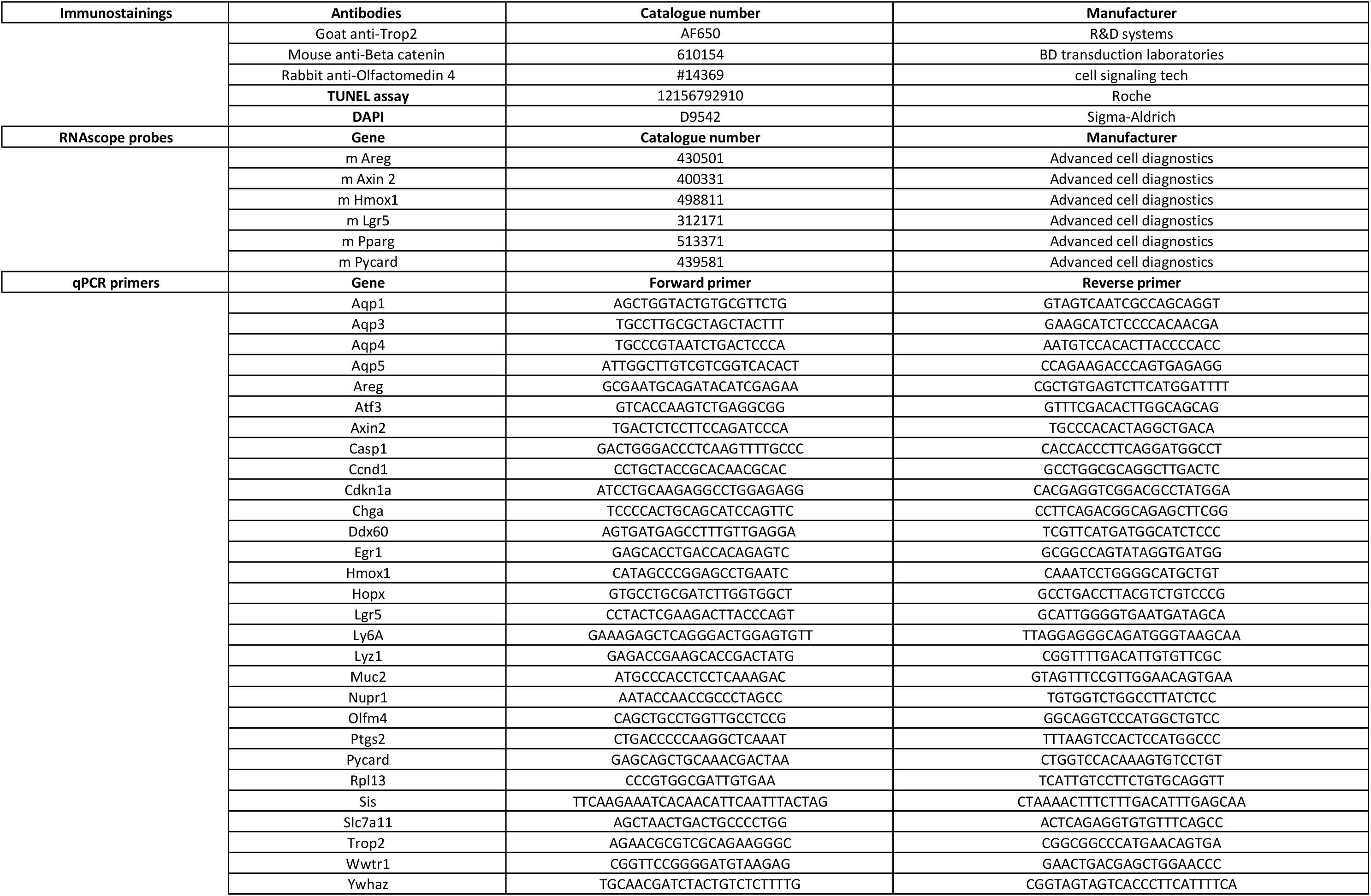
List of material used for immunostainings, in situ hybridization and qRT-PCR experiments.

### Gene expression analysis

qRT-PCR was performed on total RNA extracted from organoid cultures as reported(44). Expression levels of target genes were normalized to that of reference genes (Rpl13, Ywhaz). Table 2 lists the primers used for qPCR studies. *In situ* hybridization experiments were performed according to manufacturer instructions with the RNAscope kit (ACD-Biotechne) (probes listed in Table 2). Stained samples were visualized with a Nanozoomer digital scanner (Hamamatsu).

### RNA seq and Gene Set Enrichment Analysis (GSEA)

RNA quality was checked using a Bioanalyzer 2100 (Agilent technologies). Indexed cDNA libraries were obtained using the Ovation Solo (NuGen) or the NEBNext RNA-Seq Systems following manufacturer recommendations. The multiplexed libraries were loaded onto a NovaSeq 6000 (Illumina) using an S2 flow cell and sequences were produced using a 200 Cycle Kit. Paired-end reads were mapped against the mouse reference genome GRCm38 using STAR software to generate read alignments for each sample. Annotations Mus_musculus.GRCm38.90.gtf were obtained from https://ftp.Ensembl.org. After transcripts assembling, gene level counts were obtained using HTSeq. Differentially expressed genes were identified with EdgeR method and further analyzed using GSEA MolSig (Broad Institute)(45). Heatmaps were generated using Heatmapper(46) and common signature genes were identified using Venny 2.0 (47).

### Detection of reactive species

Detection of reactive species was performed in organoid culture supernatants 24 hours after CAP treatment, as described in (22). Nitrite concentration was measured using a colorimetric assay with the Griess reagent. Absorbance was determined at 570 nm using the iMark Microplate reader (BioRad). Global ROS were detected using the reporter 2’,7’-dichlorodihydrofluorescein diacetate (DCFH-DA, D6883 Sigma-Aldrich). The emitted fluorescence of oxidized DFC was detected at 528 nm using the Microwin software on Mithras LB940 reader (Berthold technologies).

### Statistical analysis

Statistical analyses were performed with Graph Pad Prism 5. All experimental data are expressed as mean ± s.e.m unless indicated in Figure legends. The significance of differences between groups was determined by appropriate parametric or non-parametric tests as described in the text or Figure legends.

### Data availability statement

The datasets generated and analyzed during the current study are available in the GEODATASET repository [GEO Accession GSE 178148]. Some datasets analyzed during this study were included in a published article(48) and are available in the GEODATASET repository [GSE 135362].

## ACKNOWLEDGMENTS

We are grateful to Sylvie Robine for providing the Vil1-cre/ERT2^23Syr/J^ mouse strain. We acknowledge the contribution of Maryam Marefati for tamoxifen injections and a medical writer, Sandy Field, PhD, for English language editing of this manuscript.

## CONFLICT OF INTEREST STATEMENT

The authors have nothing to disclose.

## AUTHOR CONTRIBUTION STATEMENT

AH, ML, OB, MT: study concept and design, acquisition of data, analysis and interpretation of data, statistical analysis, drafting of the ms

JD, AN, AD, FR: study concept and design, study supervision, critical revision of the ms, obtained funding

MIG: study concept and design, acquisition of data, analysis and interpretation of data, drafting of the ms, study supervision, obtained funding.

## ETHIC STATEMENTS

Animal procedures complied with the guidelines of the European Union and were approved by the local ethics committee CEBEA from Erasme-Faculty of Medicine under the accepted protocol 713N.

## FUNDING STATEMENTS

This work was supported by the “Actions de recherche concertées” (ARC, Collective Research Initiatives), the non-for-profit Association Recherche Biomédicale et Diagnostic (ARBD), and the Fondation Michel Cremer.

---

**Figure S1.**
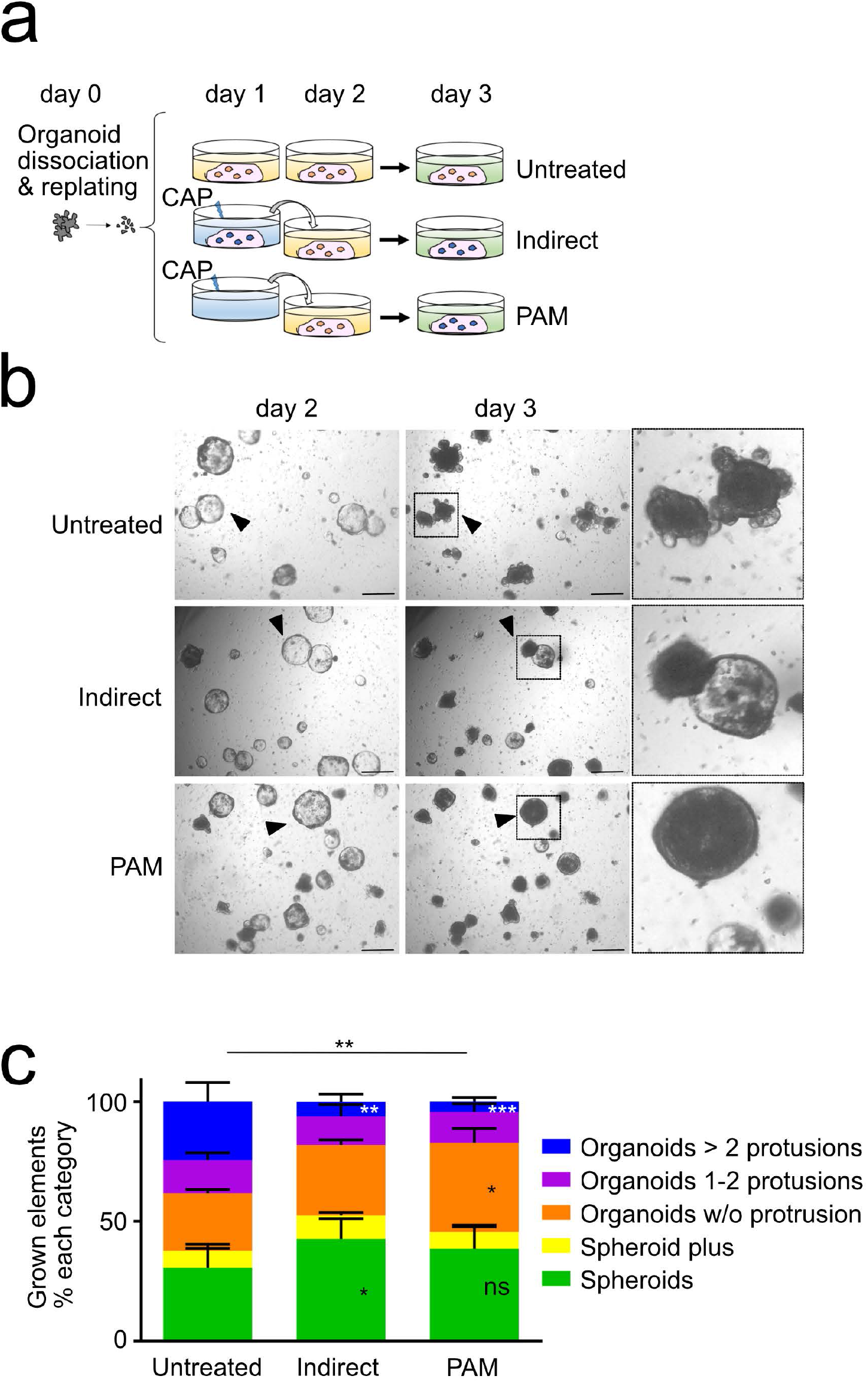
Impact of the CAP application method on organoid morphology. **a**. CAP-conditioned media and PAM generated by treatment with 50 W for 60 s at day 1 were applied directly to organoid cultures at day 2 post-replating for 24 hours (until day 3). **b**. Representative pictures of a given field showing growth of organoids at day 2 (before CAP application) and day 3 (endpoint of the experiment). Triangles show individual elements evolving as protruded organoids and spheroids in untreated and CAP-treated cultures, respectively. Right panels: insets of the pictures at day 3. Scale bars: 500 μm. **c**. Quantification of organoid complexity at day 3. An average number of 100 elements was analyzed over time per condition per organoid line (n= 4 organoid lines). Data are represented as means ± sem. Two-way ANOVA: interaction ** P< 0.01 followed by Dunnett’s multiple comparisons test: *** P< 0.001, ** P< 0.01, * P< 0.05, ns not significant (all compared to untreated).

**Figure S2.**
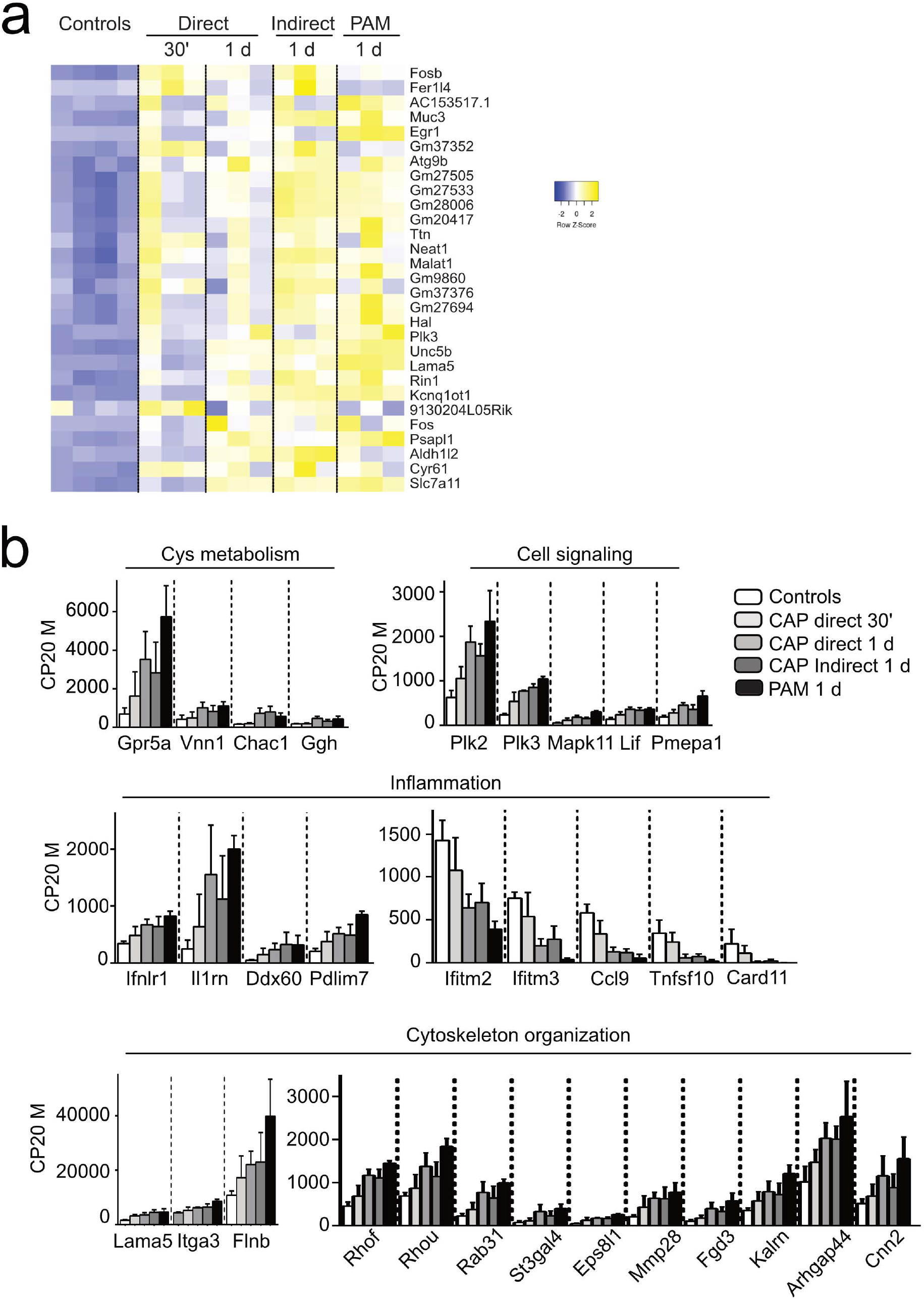
Impact of the CAP application method on global gene expression of intestinal organoids. **a**. Heatmap of the most differentially regulated genes in CAP-treated versus Untreated (Controls) organoids at the early time post-treatment (30 min). **b**. Expression levels of Cys metabolism-, cell signaling-, inflammation- and cytoskeleton organization-associated genes in the various conditions. Data are represented as means ± sd. n = 4 and 3 samples in Controls and CAP-treated conditions, respectively. CP20M: counts per kilobase of transcript per 20 million mapped reads.

**Figure S3.**
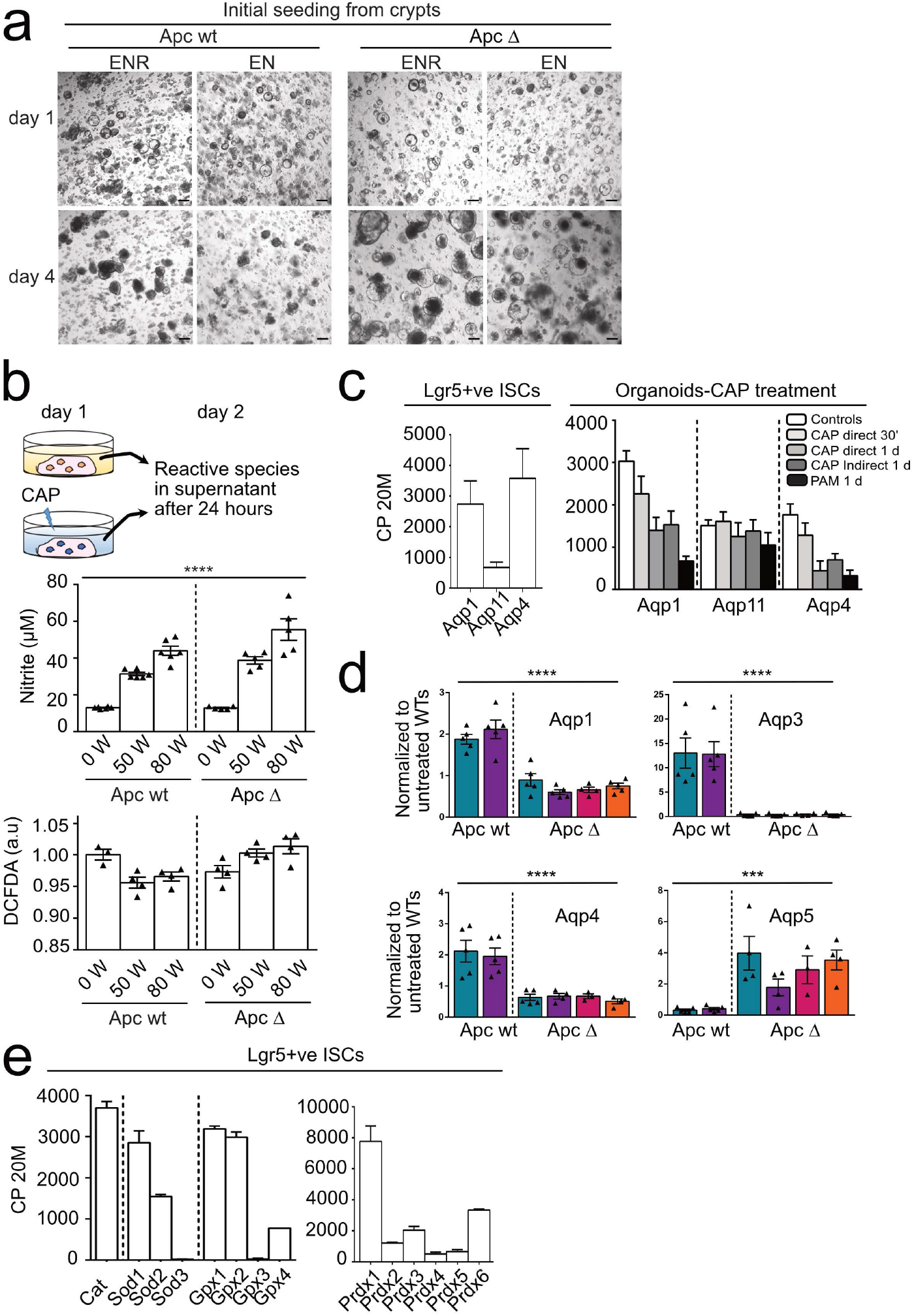
Apc deficient-derived organoids exhibit increased resistance to CAP treatment as compared to normal intestinal stem cell-derived organoids. **a**. Organoid growth of Apc wt and Apc Δ crypts upon initial seeding in culture media containing ENR (EGF, Noggin, Rspondin1) or EN (EGF, Noggin). Representative pictures of a given field showing organoid morphology at day 1 and day 4. Apc Δ organoid efficiently grow in EN conditions as compared to Apc wt organoids. Scale bars: 500 μm. **b**. Measurement of reactive species in culture supernatants of Apc wt and Apc Δ organoids 24 hours after direct CAP treatment at the indicated doses. DCFDA dye was used to measure ROS levels. A.U. Arbitrary Units. Each symbol corresponds to a given organoid line. Data are represented as means ± sem. One-way Anova test. **** P< 0.0001. **c**. Expression levels of aquaporin-encoding genes. Left panel: Aqp expression in Lgr5+ve ISC. Data were analyzed from the Gene Expression Omnibus GSE135362 dataset; Right panel: Aqp expression in control and CAP-treated organoids (this work). CP20M: counts per kilobase of transcript per 20 million mapped reads (n =2 samples). **d**. Gene expression analysis by qRT-PCR of Aquaporins in Apc wt and Apc Δ organoids. Each symbol corresponds to a given organoid line. Values are normalized to Untreated Apc wt levels. Data are represented as means ± sem. One-way Anova test. **** P< 0.0001; *** P< 0.001. **e**. Expression levels of ROS scavenger enzymes in Lgr5+ve ISC. Data were analyzed from the Gene Expression Omnibus GSE135362 dataset. CP20M: counts per kilobase of transcript per 20 million mapped reads (n=2 samples).

## Notes

### Competing Interest Statement

The authors have declared no competing interest.

### Summary of Updates

Acknowledgement section added: We acknowledge the contribution of Maryam Marefati for tamoxifen injections.

## REFERENCES

1. Yan D, Sherman JH, Keidar M. Cold atmospheric plasma, a novel promising anti-cancer treatment modality. Vol. 8, Oncotarget. Impact Journals LLC; 2017. p. 15977–95.

2. Laroussi M. Cold Plasma in Medicine and Healthcare: The New Frontier in Low Temperature Plasma Applications. Front Phys [Internet]. 2020 Mar 20 [cited 2021 Oct 8];8:74. Available from: https://www.frontiersin.org/article/10.3389/fphy.2020.00074/full

3. Dubuc A, Monsarrat P, Virard F, Merbahi N, Sarrette JP, Laurencin-Dalicieux S, et al. Use of cold-atmospheric plasma in oncology: a concise systematic review. Vol. 10, Therapeutic Advances in Medical Oncology. SAGE Publications Inc.; 2018.

4. Nakamura K, Peng Y, Utsumi F, Tanaka H, Mizuno M, Toyokuni S, et al. Novel Intraperitoneal Treatment With Non-Thermal Plasma-Activated Medium Inhibits Metastatic Potential of Ovarian Cancer Cells. Sci Rep. 2017 Dec 1;7(1).

5. Vandamme M, Robert E, Lerondel S, Sarron V, Ries D, Dozias S, et al. ROS implication in a new antitumor strategy based on non-thermal plasma. Int J Cancer. 2012 May 1;130(9):2185–94.

6. Sun Y, Lu Y, Saredy J, Wang X, Drummer IV C, Shao Y, et al. ROS systems are a new integrated network for sensing homeostasis and alarming stresses in organelle metabolic processes. Vol. 37, Redox Biology. Elsevier B.V.; 2020.

7. Bekeschus S, Liebelt G, Menz J, Berner J, Sagwal SK, Wende K, et al. Tumor cell metabolism correlates with resistance to gas plasma treatment: The evaluation of three dogmas. Free Radic Biol Med. 2021 May 1;167:12–28.

8. Perillo B, Di Donato M, Pezone A, Di Zazzo E, Giovannelli P, Galasso G, et al. ROS in cancer therapy: the bright side of the moon. Vol. 52, Experimental and Molecular Medicine. Springer Nature; 2020. p. 192–203.

9. Barker N, Van Es JH, Kuipers J, Kujala P, Van Den Born M, Cozijnsen M, et al. Identification of stem cells in small intestine and colon by marker gene Lgr5. Nature. 2007 Oct 25;449(7165):1003–7.

10. Sprangers J, Zaalberg IC, Maurice MM. Organoid-based modeling of intestinal development, regeneration, and repair. Vol. 28, Cell Death and Differentiation. Springer Nature; 2021. p. 95–107.

11. Sato T, Vries RG, Snippert HJ, Van De Wetering M, Barker N, Stange DE, et al. Single Lgr5 stem cells build crypt-villus structures in vitro without a mesenchymal niche. Nature. 2009 May 14;459(7244):262–5.

12. Van De Wetering M, Francies HE, Francis JM, Bounova G, Iorio F, Pronk A, et al. Prospective derivation of a living organoid biobank of colorectal cancer patients. Cell. 2015 May 7;161(4):933–45.

13. Roerink SF, Sasaki N, Lee-Six H, Young MD, Alexandrov LB, Behjati S, et al. Intra-tumour diversification in colorectal cancer at the single-cell level. Nature. 2018 Apr 26;556(7702):437–62.

14. Yao Y, Xu X, Yang L, Zhu J, Wan J, Shen L, et al. Patient-Derived Organoids Predict Chemoradiation Responses of Locally Advanced Rectal Cancer. Cell Stem Cell. 2020 Jan 2;26(1):17–26.e6.

15. Tuveson D, Clevers H. Cancer modeling meets human organoid technology. Vol. 364, Science. American Association for the Advancement of Science; 2019. p. 952–5.

16. Driehuis E, Kretzschmar K, Clevers H. Establishment of patient-derived cancer organoids for drug-screening applications. Nat Protoc. 2020 Oct 1;15(10):3380–409.

17. Muñoz J, Stange DE, Schepers AG, Van De Wetering M, Koo BK, Itzkovitz S, et al. The Lgr5 intestinal stem cell signature: Robust expression of proposed quiescent ′ +4′ cell markers. EMBO J. 2012 Jul 18;31(14):3079–91.

18. Fernandez Vallone V, Leprovots M, Strollo S, Vasile G, Lefort A, Libert F, et al. Trop2 marks transient gastric fetal epithelium and adult regenerating cells after epithelial damage. Development [Internet]. 2016 May 1 [cited 2019 Sep 10];143(9):1452–63. Available from: http://dev.biologists.org/lookup/doi/10.1242/dev.131490

19. Ghafouri-Fard S, Shoorei H, Taheri M. Non-coding RNAs are involved in the response to oxidative stress. Vol. 127, Biomedicine and Pharmacotherapy. Elsevier Masson SAS; 2020.

20. Korbecki J, Bobiński R, Dutka M. Self-regulation of the inflammatory response by peroxisome proliferator-activated receptors. Vol. 68, Inflammation Research. Birkhauser Verlag AG; 2019.

21. Yan D, Talbot A, Nourmohammadi N, Cheng X, Canady J, Sherman J, et al. Principles of using Cold Atmospheric Plasma Stimulated Media for Cancer Treatment. Sci Rep. 2015 Dec 17;5.

22. Tornin J, Labay C, Tampieri F, Ginebra MP, Canal C. Evaluation of the effects of cold atmospheric plasma and plasma-treated liquids in cancer cell cultures. Vol. 16, Nature Protocols. Nature Research; 2021. p. 2826–50.

23. Guarino VA, Oldham WM, Loscalzo J, Zhang YY. Reaction rate of pyruvate and hydrogen peroxide: assessing antioxidant capacity of pyruvate under biological conditions. Sci Rep. 2019 Dec 1;9(1).

24. Yang X, Chen G, Yu KN, Yang M, Peng S, Ma J, et al. Cold atmospheric plasma induces GSDME-dependent pyroptotic signaling pathway via ROS generation in tumor cells. Cell Death Dis. 2020 Apr 1;11(4).

25. Nishi K, Iwaihara Y, Tsunoda T, Doi K, Sakata T, Shirasawa S, et al. ROS-induced cleavage of NHLRC2 by caspase-8 leads to apoptotic cell death in the HCT116 human colon cancer cell line article. Cell Death Dis. 2017 Dec 1;8(12).

26. Beyer RE. The role of ascorbate in antioxidant protection of biomembranes: Interaction with vitamin E and coenzyme Q. J Bioenerg Biomembr. 1994 Aug;26(4):349–58.

27. Yui S, Azzolin L, Maimets M, Pedersen MT, Fordham RP, Hansen SL, et al. YAP/TAZ-Dependent Reprogramming of Colonic Epithelium Links ECM Remodeling to Tissue Regeneration. Cell Stem Cell. 2018 Jan 4;22(1):35–49.e7.

28. Ayyaz A, Kumar S, Sangiorgi B, Ghoshal B, Gosio J, Ouladan S, et al. Single-cell transcriptomes of the regenerating intestine reveal a revival stem cell. Nature [Internet]. 2019; Available from: http://dx.doi.org/10.1038/s41586-019-1154-y

29. Nusse YM, Savage AK, Marangoni P, Rosendahl-Huber AKM, Landman TA, De Sauvage FJ, et al. Parasitic helminths induce fetal-like reversion in the intestinal stem cell niche. Nature. 2018 Jul 5;559(7712):109–13.

30. Wang Y, Chiang IL, Ohara TE, Fujii S, Cheng J, Muegge BD, et al. Long-Term Culture Captures Injury-Repair Cycles of Colonic Stem Cells. Cell [Internet]. 2019;179(5):1144–1159.e15. Available from: https://doi.org/10.1016/j.cell.2019.10.015

31. Eggers B, Marciniak J, Deschner J, Stope MB, Mustea A, Kramer FJ, et al. Cold atmospheric plasma promotes regeneration‐associated cell functions of murine cementoblasts in vitro. Int J Mol Sci. 2021 May 2;22(10).

32. Kwong M, Kan YW, Chan JY. The CNC basic leucine zipper factor, Nrf1, is essential for cell survival in response to oxidative stress-inducing agents. Role for Nrf1 in γ-gcs(L) and gss expression in mouse fibroblasts. J Biol Chem. 1999 Dec 24;274(52):37491–8.

33. Tanigawa S, Lee CH, Lin CS, Ku CC, Hasegawa H, Qin S, et al. Jun dimerization protein 2 is a critical component of the Nrf2/MafK complex regulating the response to ROS homeostasis. Cell Death Dis. 2013;4(11).

34. Tabuchi Y, Uchiyama H, Zhao QL, Yunoki T, Andocs G, Nojima N, et al. Effects of nitrogen on the apoptosis of and changes in gene expression in human lymphoma U937 cells exposed to argon-based cold atmospheric pressure plasma. Int J Mol Med. 2016 Jun 1;37(6):1706–14.

35. Kou L, Jiang X, Huang H, Lin X, Zhang Y, Yao Q, et al. The role of transporters in cancer redox homeostasis and cross-talk with nanomedicines. Vol. 15, Asian Journal of Pharmaceutical Sciences. Shenyang Pharmaceutical University; 2020. p. 145–57.

36. Semmler ML, Bekeschus S, Schäfer M, Bernhardt T, Fischer T, Witzke K, et al. Molecular mechanisms of the efficacy of cold atmospheric pressure plasma (CAP) in cancer treatment [Internet]. Vol. 12, Cancers. MDPI AG; 2020 [cited 2020 Feb 21]. Available from: http://www.ncbi.nlm.nih.gov/pubmed/31979114

37. Fodde R. The APC gene in colorectal cancer. Eur J Cancer. 2002;38(7):867–71.

38. Malki A, Elruz RA, Gupta I, Allouch A, Vranic S, Al Moustafa AE. Molecular mechanisms of colon cancer progression and metastasis: Recent insights and advancements. Vol. 22, International Journal of Molecular Sciences. MDPI AG; 2021. p. 1–24.

39. El Marjou F, Janssen KP, Chang BHJ, Li M, Hindie V, Chan L, et al. Tissue-specific and inducible Cre-mediated recombination in the gut epithelium. Genesis. 2004 Jul;39(3):186–93.

40. Robanus-Maandag EC, Koelink PJ, Breukel C, Salvatori DCF, Jagmohan-Changur SC, Bosch CAJ, et al. A new conditional Apc-mutant mouse model for colorectal cancer. Carcinogenesis. 2010 Feb 22;31(5):946–52.

41. Vallone V, Leprovots M, Vassart G, Garcia M-I. Ex vivo Culture of Fetal Mouse Gastric Epithelial Progenitors. BIO-PROTOCOL. 2017;7(1).

42. Bastin O, Thulliez M, Servais J, Nonclercq A, Delchambre A, Hadefi A, et al. Optical and electrical characteristics of an endoscopic DBD plasma jet. Plasma Med. 2020;10(2):71–90.

43. Garcia MI, Ghiani M, Lefort A, Libert F, Strollo S, Vassart G. LGR5 deficiency deregulates Wnt signaling and leads to precocious Paneth cell differentiation in the fetal intestine. Dev Biol. 2009 Jul 1;331(1):58–67.

44. Mustata RC, Van Loy T, Lefort A, Libert F, Strollo S, Vassart G, et al. Lgr4 is required for Paneth cell differentiation and maintenance of intestinal stem cells ex vivo. EMBO Rep. 2011;12(6):558–64.

45. Subramanian A, Tamayo P, Mootha VK, Mukherjee S, Ebert BL, Gillette MA, et al. Gene set enrichment analysis: A knowledge-based approach for interpreting genome-wide expression profiles. Proc Natl Acad Sci U S A. 2005 Oct 25;102(43):15545–50.

46. Babicki S, Arndt D, Marcu A, Liang Y, Grant JR, Maciejewski A, et al. Heatmapper: web-enabled heat mapping for all. Nucleic Acids Res. 2016 Jul 8;44(W1):W147–53.

47. Oliveros JC (2007-2015). Venny 2.1.0 [Internet]. An interactive tool for comparing lists with Venn’s diagrams. [cited 2021 Sep 25]. Available from: https://bioinfogp.cnb.csic.es/tools/venny/index.html

48. Fernandez Vallone V, Leprovots M, Ribatallada‐Soriano D, Gerbier R, Lefort A, Libert F, et al. LGR 5 controls extracellular matrix production by stem cells in the developing intestine. EMBO Rep. 2020 Jul 3;21(7).

